# The exported chaperone PfHsp70x is dispensable for the *Plasmodium falciparum* intraerythrocytic lifecycle

**DOI:** 10.1101/113365

**Authors:** David W. Cobb, Anat Florentin, Manuel A. Fierro, Michelle Krakowiak, Julie M. Moore, Vasant Muralidharan

## Abstract

Export of parasite proteins into the host erythrocyte is essential for survival of *Plasmodium falciparum* during its asexual lifecycle. While several studies described key factors within the parasite that are involved in protein export, the mechanisms employed to traffic exported proteins within the host cell are currently unknown. Members of the Hsp70 family of chaperones, together with their Hsp40 co-chaperones, facilitate protein trafficking in other organisms, and are thus likely used by *P. falciparum* in the trafficking of its exported proteins. A large group of Hsp40 proteins is encoded by the parasite and exported to the host cell, but only one Hsp70, PfHsp70x, is exported with them.

PfHsp70x is absent from most *Plasmodium* species and is found only in *P. falciparum* and closely-related species that infect Apes. Herein, we have utilized CRISPR/Cas9 genome editing in *P. falciparum* to investigate the essentiality of PfHsp70x. We show that parasitic growth was unaffected by knockdown of PfHsp70x using both the DHFR-based Destabilization Domain and the *glmS* ribozyme system. Similarly, a complete gene knockout of PfHsp70x did not affect the ability of *P. falciparum* to proceed through its intraerythrocytic lifecycle. The effect of PfHsp70x knockdown/knockout on the export of proteins to the host RBC, including the critical virulence factor PfEMP1, was tested and we found that this process was unaffected. These data show that although PfHsp70x is the sole exported Hsp70, it is not essential for the asexual development of *P. falciparum*.

**Importance:** Half of the world’s population lives at risk for malaria. The intraerythrocytic lifecycle of *Plasmodium* spp. is responsible for clinical manifestations of malaria; therefore, knowledge of the parasite’s ability to survive within the erythrocyte is needed to combat the deadliest agent of malaria, *P. falciparum*. An outstanding question in the field is how *P. falciparum* undertakes the essential process of trafficking its proteins within the host cell. In most organisms, chaperones such as Hsp70 are employed in protein trafficking. Of the human-disease causing Plasmodium species, the chaperone PfHsp70x is unique to *P. falciparum,* and it is the only parasite protein of its kind exported to the host (1). This has placed PfHsp70x as an ideal target to inhibit protein trafficking and kill the parasite. However, we show that PfHsp70x is not required for export of parasite effectors nor is it essential for parasite survival inside of the RBC.

## Introduction

Malaria is a profound killer worldwide. In 2015, 214 million cases of malaria resulted in 438,000 deaths, largely in Africa and Asia (2). Within malaria endemic countries, the disease targets the most vulnerable of the population, including children under five and pregnant women (2). The disease is caused by infection with eukaryotic parasites from the genus *Plasmodium*, but it is one species*—P. falciparum*—that is responsible for most of the malaria associated mortality. The clinical manifestations of malaria range from fever, headache, and muscle pains, to severe anemia, coma, and respiratory distress (3). All of these symptoms are direct consequences of asexual replication of the parasite within the human red blood cell (RBC)(4). During this cycle of replication, *P. falciparum* invades the RBC and dramatically transforms its morphology and physiology. Alterations to the RBC include increased permeability, loss of cell deformability, and introduction of virulence-associated knobs at the RBC membrane (5, 6).

Remodeling of the RBC requires export of hundreds of parasite proteins into the host cell, a feat involving protein trafficking through multiple compartments before arriving at their final destinations in the host. The first phase of the journey begins in the parasite endoplasmic reticulum (ER). Many exported proteins contain an N-terminal signal motif termed the Host Targeting Signal or *Plasmodium* Export Element (PEXEL) (6, 7). A key step in the export of PEXEL-containing proteins is cleavage of the motif by the ER-resident aspartyl protease Plasmepsin V (8–10). A sub-group of exported proteins called PEXEL-Negative Exported Proteins (PNEPs) lack the motif, but their N-terminus is similarily necessary for export (12, 13). Aside from Plasmepsin V processing of PEXEL, mechanisms underlying the selection of host-destined proteins for exit from the ER remain unclear. Nonetheless, PEXEL-proteins and PNEPs continue their journey through the parasite’s secretory pathway and are delivered to the parasitophorous vacuole (PV), a membranous structure within which the parasite resides. Previous studies have shown that proteins cross the parasitophorous vacuole membrane (PVM) through the *Plasmodium* Translocon of Exported Proteins (PTEX) (14–16). Once they are on the other side of the PVM, all classes of proteins need to refold and find their specific subcellular localization, whether it is in the host cytoplasm, the host membrane, or parasite-induced structures such as knobs or Maurer’s clefts. It is completely unknown how hundreds of proteins, within a short time period, cross through PTEX, refold to regain structure and function, and find their final destination in the host.

The process of protein export is essential for *P. falciparum* survival in the RBC, as blockage of protein export—whether at the parasite ER or at the PVM—results in parasite death. In the ER, overexpression of catalytically dead Plasmepsin V (PMV) results in impaired parasite growth, and inhibition of PMV with a PEXEL-mimetic impairs protein export and kills parasites during the transition to the trophozoite stage (10, 17, 18). Similarly, *P. falciparum* parasites are sensitive to interference of trafficking across the PVM. Conditional knockdown of PTEX components blocks protein export and kills the parasites (19, 20). As the parasites are susceptible to inhibition of trafficking in the ER and PV, interference in the trafficking process within the host may similarly impair parasite growth. The mechanisms of protein trafficking inside of the host cell remain unknown, but identification of essential components of this process will provide valuable targets for drug discovery programs.

Molecular chaperones are likely candidates in the search for key export and trafficking components. Indeed, PfHsp101 is an essential component of PTEX, and its inhibition results in accumulation of exported proteins within the PV (19). Furthermore, several parasite Hsp40s are exported to the RBC, but their function there is unknown (21). In other organisms Hsp40s serve as co-chaperones for Hsp70s, but in contrast to the large number of exported Hsp40s, PfHsp70x (PF3D7_0831700) is the only parasite-encoded Hsp70 that is exported to host cell (1, 22). This chaperone is found only in *P. falciparum* and closely related species that cause malaria in apes such as *P. reichenowi,* but not in other *Plasmodium* species that infect humans, such as *P. vivax* or *P. knowlesi* (1). Within the *P. falciparum* infected RBC, PfHsp70x is localized to the PV and the host, where it associates with PfHsp40s in mobile structures termed J-dots (1). Given its status as sole exported Hsp70, we hypothesized that PfHsp70x is central to protein trafficking in the host cell, and thus essential to parasite viability. Indeed, studies focused on PTEX interactions have found PfHsp70x associated with the translocon, and it has been shown to co-localize with the critical virulence protein PfEMP1 during its trafficking (1, 23, 24).

In this study, we took advantage of various genetic techniques to show that PfHsp70x is non-essential for protein export and parasite growth. We have used the DHFR-based destabilizing domain that has previously been used to inhibit chaperone function (19, 25). In addition, we have used the *glmS*-ribozyme system that inhibits translation via mRNA degradation (26). Mutants for both knockdown methods were successfully generated, but knockdown had no impact on parasite growth or protein export, including no discernible difference in the export of PfEMP1. To confirm that the lack of a phenotype was not due to incomplete knockdown, we used CRISPR/Cas9 technology to generate a complete knockout of the PfHsp70x gene and found no defects in parasite proliferation or export. Our data demonstrate that PfHsp70x is not required for protein export to the host RBC and not essential for the intraerythrocytic lifecycle of *P. falciparum*.

## Results

### Conditional mutants of PfHsp70x

Previous work has shown that the DHFR-based destabilization domain (DDD) fusions can lead to the inhibition of protein-protein interactions (19, 25) or degradation of the DDD-tagged proteins (27–29). In the presence of the stabilizing ligand trimethoprim (TMP), the DDD is folded and the chaperone functions normally. However, upon TMP removal the DDD is unfolded and binds to its attached chaperone intramolecularly, thereby blocking interactions with the chaperone’s client proteins and inhibiting normal chaperone function (**Fig. S1A**). Relying on single-crossover homologous recombination, the *pfhsp70x* gene was modified with a triple-HA tag and the DDD, and integration at the *pfhsp70x* locus was confirmed via Southern blot analysis (**Fig. S1A, B**). Consistent with the auto-inhibitory model of chaperone-DDD action, western blot analysis of parasite lysates following TMP removal showed that PfHsp70x protein levels remain consistent over time (**Fig. S1C**). Isolation of the host cell cytoplasm using saponin lysis revealed that PfHsp70x-DDD is exported to the host cell (**Fig. S1C**). Moreover, the persistence of PfHsp70x in the supernatant following TMP removal indicated that PfHsp70x is exported to the host cell even in its putative inhibited form. To assess the role of PfHsp70x in parasite proliferation, we removed TMP and measured asexual growth over a course of several days and at least two replication cycles. We found that the absence of TMP had no effect on parasite proliferation (**Fig. S2A**). It was previously reported that PfHsp70x, together with several other exported chaperones, localizes to specific punctate structures in the host cell termed J-dots. To test the effect of DDD-based inhibition on PfHsp70x localization, we performed immunofluorescence assays and found that PfHsp70x-DDD is trafficked to the expected punctate structures within the host cell, regardless of TMP presence (**Fig. S2B**). These data suggest that unlike other chaperones, PfHsp70x activity was unaffected by the DDD fusion or that inhibition of PfHsp70x using the DDD system does not affect the asexual life cycle of the parasite. We therefore utilized alternative methods to reduce PfHsp70x protein levels in the parasite.

Next, we sought to conditionally knockdown PfHsp70x at the mRNA level using the *glmS* ribozyme (26). In this system, the *glmS* ribozyme sequence is inserted into the 3’ end of the genomic locus of a gene and is transcribed with the gene as one mRNA. Addition of the small molecule glucosamine (GlcN) activates the *glmS* ribozyme, which cleaves itself from the mRNA, disconnecting the transcript from its polyA tail and leading to its degradation (**Fig. 1A**). Using CRISPR/Cas9 genome engineering, we appended a triple HA tag to the C-terminus of PfHsp70x followed by the *glmS* ribozyme (**Fig. 1A**) (30). A second cell line was generated in which the *pfhsp70x* locus was tagged with a mutant version of the ribozyme—termed *M9*—which is unresponsive to GlcN and serves as a control during GlcN treatment (26). Following transfection and drug selection, PfHsp70x*-glmS* and PfHsp70x*-M9* clones were isolated via limiting dilution. PCR analysis revealed the correct integration of the tag and ribozyme into the *pfhsp70x* gene in all clonal parasite lines (**Fig. 1B**). Additionally, immunofluorescence assays confirmed that PfHsp70x*-glmS* is exported to the host cytoplasm, where it is found, as before, in punctate structures that are distinct from Maurer’s clefts, suggestive of J-dot localization (**Fig. 1C**).

**Fig. 1.**
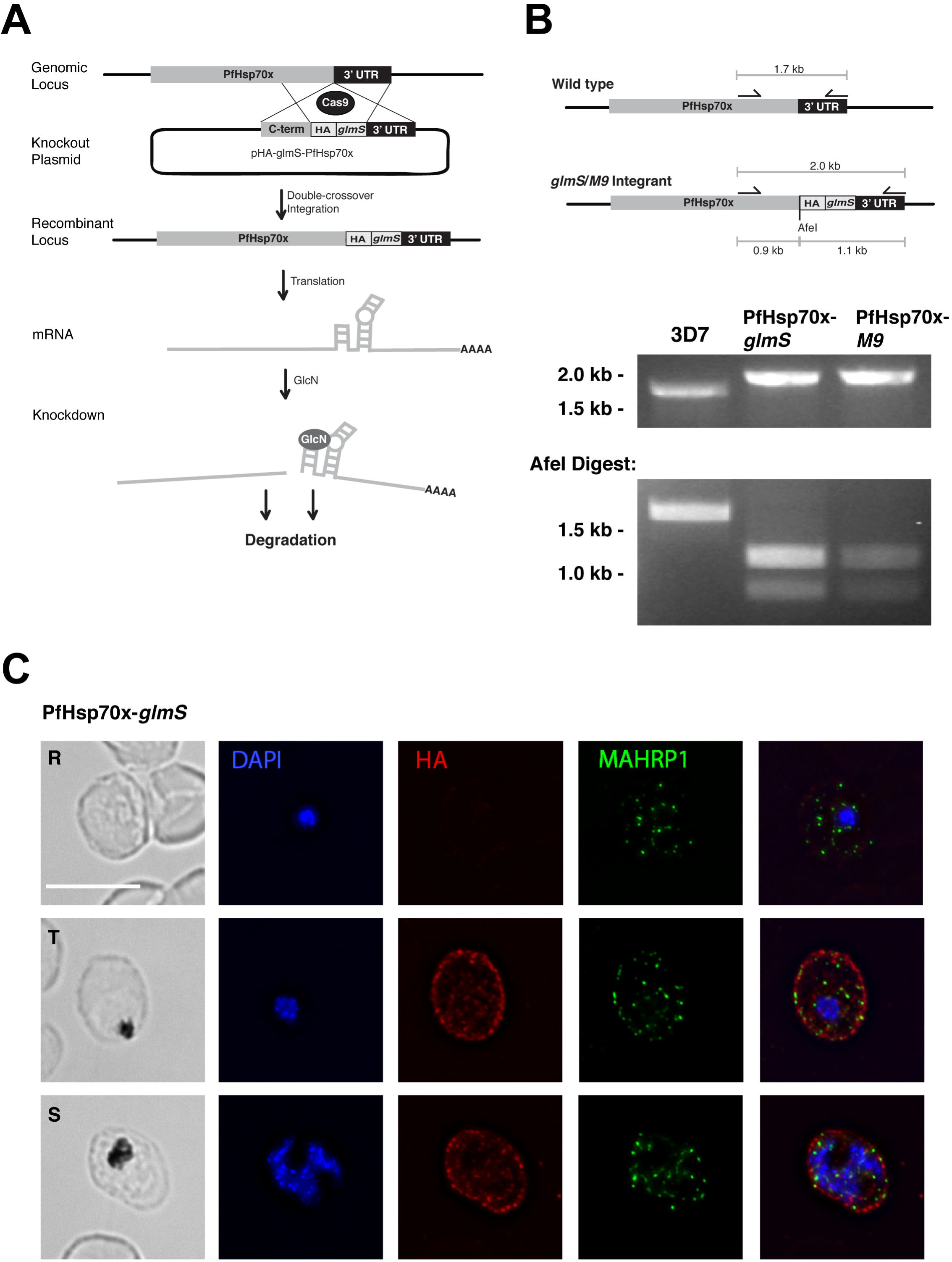
CRISPR/Cas9-mediated integration of HA-*glmS*/*M9* at PfHsp70x locus. (A) Diagram showing integration of 3xHA-ribozyme sequence and GlcN-induced degradation of mRNA. Cas9 introduces a double-stranded break at the beginning of the 3’ UTR of the *pfhsp70x* locus. The repair plasmid provides homology regions for double-crossover homologous recombination, introducing a 3xHA tag and the ribozyme sequence. Following translation and addition of GlcN, the PfHsp70x-*glmS* mRNA is cleaved by the ribozyme and is subject to degradation. (B) PCR test confirming integration at the PfHsp70x locus. DNA was purified from transfected, cloned parasites and primers were used to amplify the region between the C-terminus and 3’UTR of *pfhsp70x*. The PCR products were digested with AfeI, further confirming integration. (C) IFA showing export of HA-tagged PfHsp70x. Asynchronous PfHsp70x-*glmS* parasites were fixed with acetone and stained with specific antibodies. Images from left to right are phase, DAPI (parasite nucleus, blue), anti-HA (red), anti-MAHRP1 (green), and fluorescence merge. Scale bar represents 5 μm. (R) rings, (T) trophs and (S) schizont.

Next, we tested the effect of reducing PfHsp70x levels on intraerythrocytic growth. To ensure that insertion of the ribozyme itself does not interfere with normal asexual growth, PfHsp70x*-glmS*, PfHsp70x*-M9*, and the parental line (3D7) were grown in the absence of GlcN. Indeed, we found that in the absence of GlcN, growth of both the *glmS* and *M9* cell lines was comparable to 3D7 (**Fig. 2A**). Next, PfHsp70x*-glmS* and PfHsp70x-*M9* were cultured with GlcN and parasitemia was measured via flow cytometry. The growth of PfHsp70x*-glmS* and PfHsp70x*-M9* was unaffected by treatment with 5 mM and 10 mM GlcN (**Fig. 2B, C**). To confirm that PfHsp70x protein level is reduced in response to GlcN, schizont-stage parasites from the *glmS* and *M9* cell lines were Percoll-purified and whole parasite lysates were used for western blotting. Using anti-HA antibody we found that treatment with GlcN reduced protein levels in PfHsp70x*-glmS* but did not affect protein levels in PfHsp70x*-M9* (**Fig. 2D**). Together, these data show that we can efficiently reduce PfHsp70x levels using the *glmS* ribozyme but this has no affect on the asexual growth of the parasite within the RBC.

**Fig. 2.**
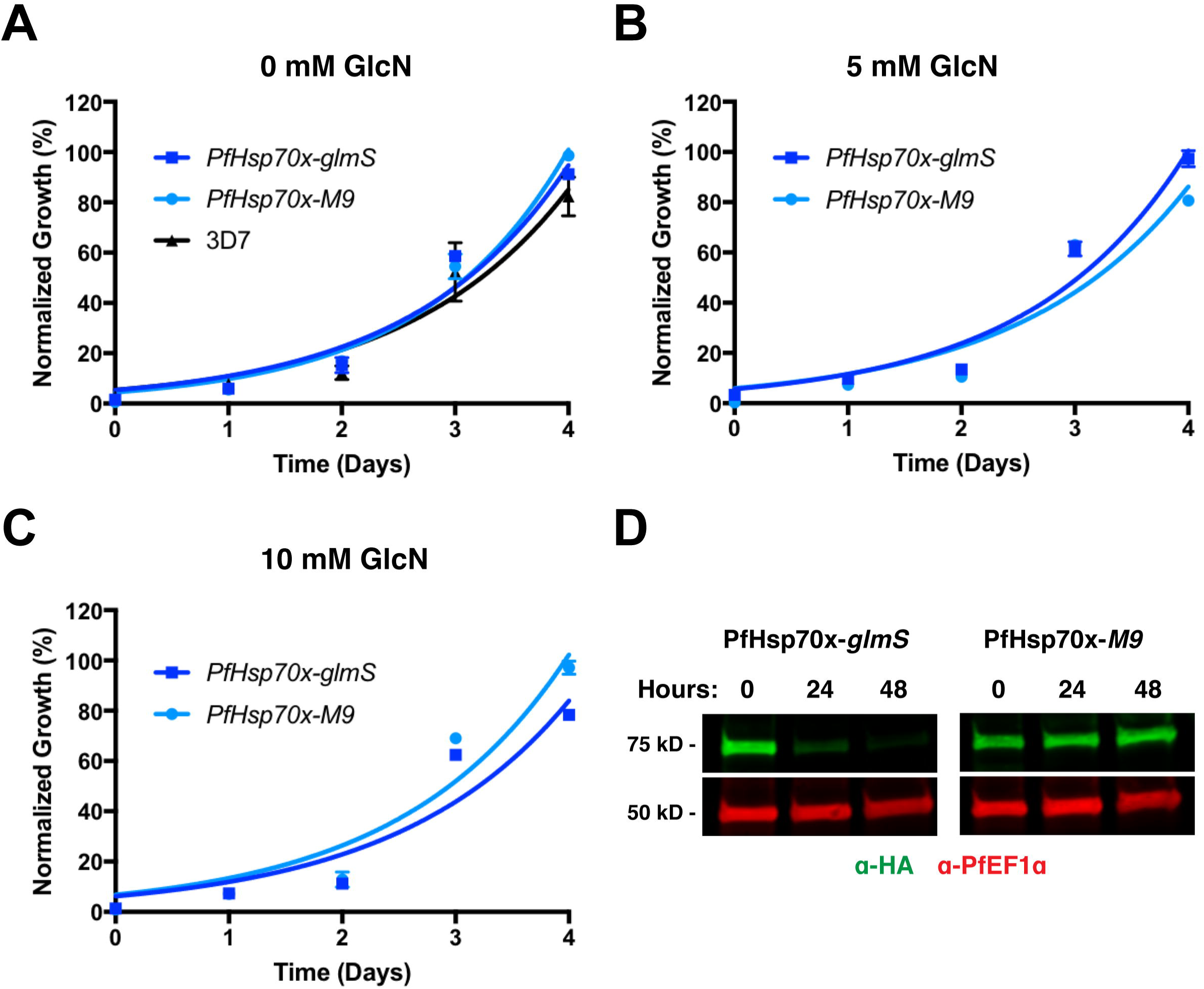
GlcN-induced knockdown of PfHsp70x does not affect intraerythrocytic growth. (A) PfHsp70x-*glmS*, PfHsp70x-*M9*, and 3D7 (parental line) were seeded at equal parasitemia in triplicate and grown in normal culturing media (complete RPMI). Parasitemia was measured every 24 hours using flow cytometry. Data are fit to an exponential growth equation and are represented as mean ± S.E.M. (n=3). (B) and (C) PfHsp70x-*glmS* and PfHsp70x-*M9* parasites were seeded at equal parasitemia in triplicate. Cultures were grown in the presence of either 5 mM (B) or 10 mM (C) GlcN. Parasitemia was measured every 24 hours using flow cytometry. Data are fit to an exponential growth equation and are represented as mean ± S.E.M. (n=3). (D) PfHsp70x-*glmS* and PfHsp70x-*M9* parasites were grown in the presence of 7.5 mM GlcN. Schizont-stage parasites were Percoll-purified every 24 hours, and whole parasite lysates were used for western blot analysis. Membrane was probed with anti-HA, and anti-PfEF1α (loading control).

### Protein export is unimpaired in PfHsp70x-knockdown parasites

Although parasite growth was unaffected by PfHsp70x knockdown, we reasoned that it could nonetheless play a role in export of proteins to the host cell. In particular, we hypothesized that PfHsp70x is needed for the export of proteins known to mediate virulence of *P. falciparum* infection, as trafficking defects of these proteins would not manifest as arrest of the asexual lifecycle (21). Using immunofluorescence, we examined localization of specific virulence-associated proteins in PfHsp70x-*M9* and PfHsp70x-*glmS* parasites after 72 hours of growth in GlcN-supplemented medium. First, the localization of the PEXEL-containing PfFIKK4.2, an exported kinase associated with knob formation and infected RBC rigidity, is unchanged in control versus PfHsp70x-knockdown parasites **(Fig. 3A)** (31). Next, we examined the localization of the PEXEL-containing protein KAHRP, which is essential for the formation of knobs on the surface of infected RBCs (32). Export of this protein was not inhibited in PfHsp70x-knockdown parasites **(Fig. 3B)**. Finally, we determined the localization of the PNEP MAHRP1, which has been implicated in the presentation of antigenically variant proteins, including PfEMP1, at the RBC surface, and we found that its export is not impaired by the knockdown of PfHsp70x **(Fig. 3C)** (33). As demonstrated by HA-staining in western blot and IFA (**Fig. 2D**, **Fig. 3**), PfHsp70x is reduced, but not completely ablated, using the *glmS* ribozyme. We reasoned that the reduced level of PfHsp70x that is produced during GlcN treatment could be sufficient for parasite survival, and therefore endeavored next to knockout *pfhsp70x*.

**Fig. 3.**
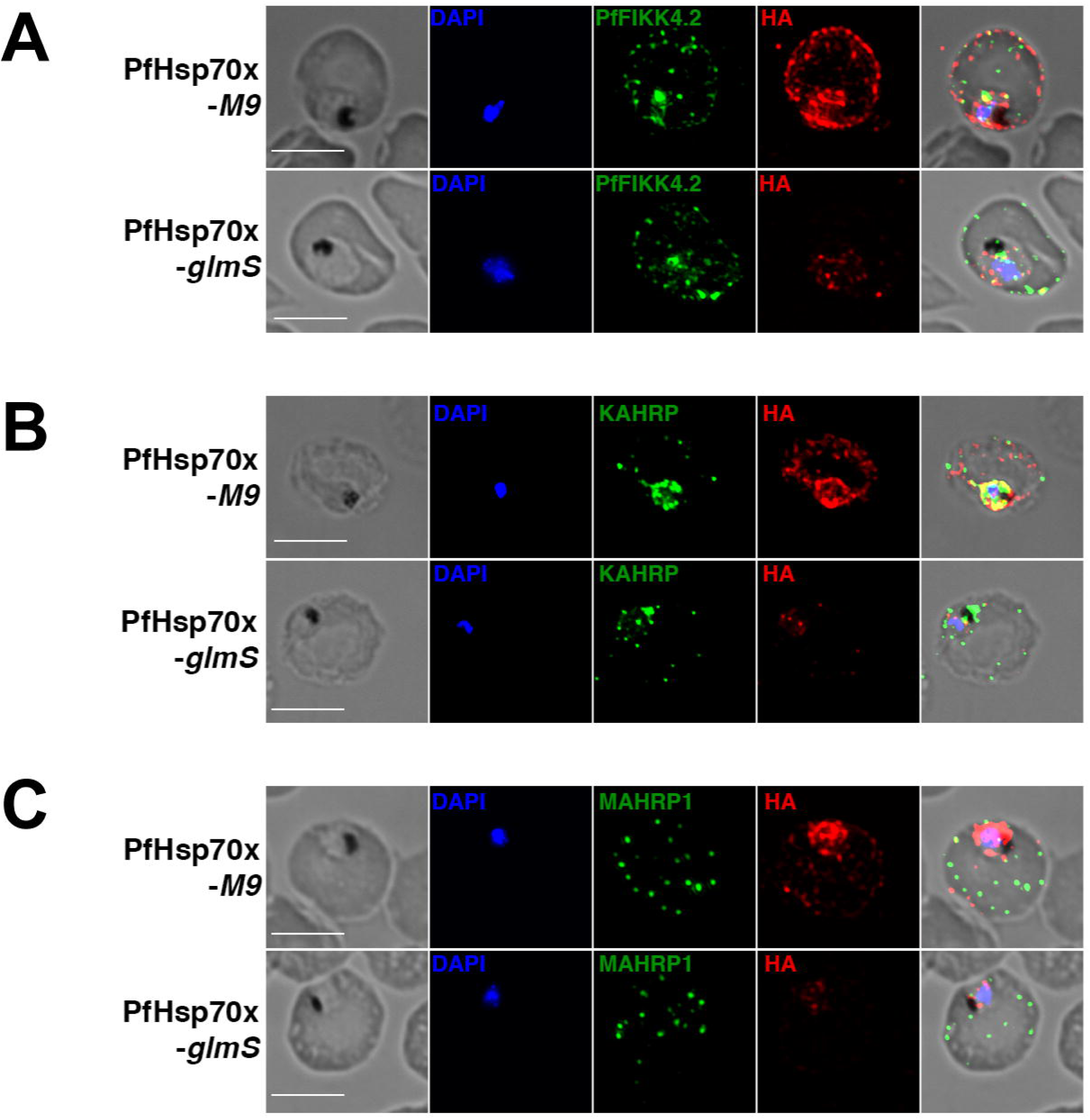
PfHsp70x knockdown does not inhibit export of virulence-associated proteins. Asynchronous PfHsp70x-*M9* and PfHsp70x-*glmS* parasites were fixed with acetone (PfFIKK4.2 and MAHRP1) or paraformaldehyde (KAHRP) and stained with antibodies against (A) PfFIKK4.2, (B) KAHRP, or (C) MAHRP1. DAPI used to mark parasite cell nucleus. Scale bar represents 5μm. Images from left to right are phase, DAPI (blue), anti-exported protein (green), anti-HA (red), and fluorescence and phase merge. Representative images shown.

### Knockout of *pfhsp70x* does not affect parasite growth

We utilized two different conditional knockdown systems to modify the PfHsp70x locus, but these approaches were insufficient to produce a growth defect in the parasites. Therefore, we sought to definitively test the essentiality of PfHsp70x via complete genomic knockout (termed PfHsp70x-KO). To this end, we employed CRISPR/Cas9 to interrupt the PfHsp70x ORF by inserting a human dihydrofolate reductase (*hdhfr*) drug resistance cassette (**Fig. 4A**). Following transfection and selection with WR99210, PfHsp70x-KO parasites were cloned via limiting dilution. Southern blot analysis of genomic DNA isolated from the parental line and independent clones showed that the *hdhfr* cassette was inserted into the *pfhsp70x* gene via homology directed repair (**Fig. 4B**). To verify that the null mutants do not express PfHsp70x, schizont-stage parasites from two independent knockout clones and the parental line were Percoll-purified, and whole parasite lysates were used for western blotting. Probing with anti-PfHsp70x shows that the knockout clones do not express PfHsp70x (**Fig. 4C**). Intraerythrocytic growth of the PfHsp70x-KO clones was monitored over two replication cycles. In agreement with the lack of any growth phenotype in the conditional knockdown parasite lines, the PfHsp70x-KO parasites displayed *wild-type* level of proliferation in erythrocytes (**Fig. 4D**). Finally, we measured the susceptibility of PfHsp70x-KO clones to heat shock stress by monitoring their growth after a heat shock (**Fig. S3)**. These data show that the PfHsp70x-KO parasites are able to deal with heat shock just as well as the wild type parasites (**Fig. S3**). The normal growth in the complete absence of PfHsp70x expression conclusively demonstrates that PfHsp70x activity is not essential for the asexual growth of the parasite within the RBC.

**Fig. 4.**
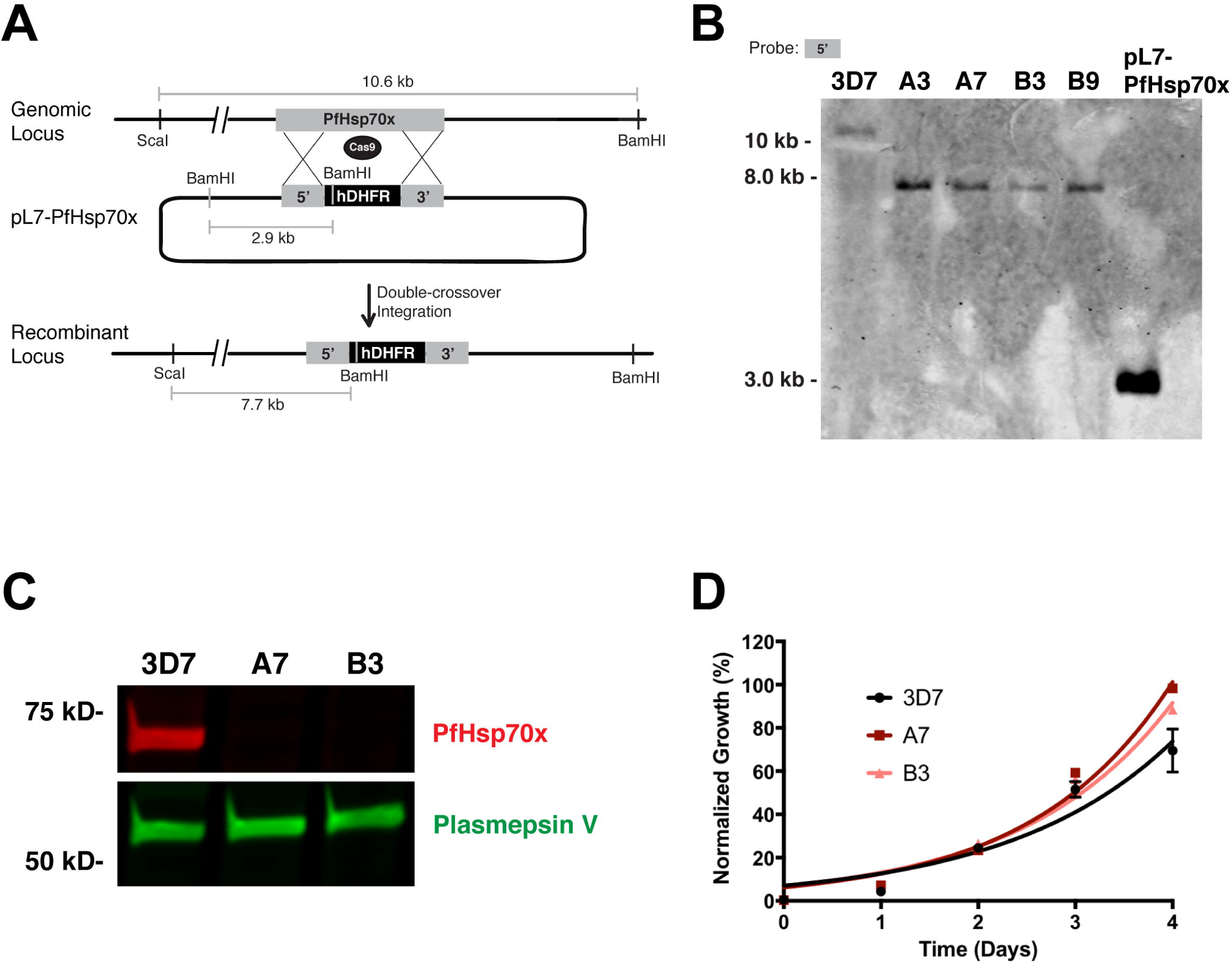
Knockout of *pfhsp70x* does not affect intraerythrocytic growth. (A) Schematic showing interruption of PfHsp70x ORF with *hDHFR* cassette. Cas9-mediated double-stranded break in the *pfhsp70x* ORF is repaired using homology regions on the template plasmid while inserting an *hDHFR* cassette into the locus. (B) Southern blot analysis confirming knockout of PfHsp70x. Genomic DNA from independent knockout clones (A3, A7, B3, and B9) was isolated and digested with BamHI and ScaI. Membrane was hybridized with a biotin-labeled probe complementary to the first 800 base pairs of the *pfhsp70x* ORF. (C) Western blot analysis demonstrating loss of PfHsp70x protein expression in independent knockout clones. Schizont-stage parasites were Percoll-purified, and whole cell lysate was used for analysis. Membrane was probed with antibodies raised against PfHsp70x, and against Plasmepsin V as a loading control. (D) Parental lines and independent PfHsp70x-KO clones (A7 and B3) were seeded at equal parasitemia in triplicate. Parasitemia was measured every 24 hours using flow cytometry. Data are fit to an exponential growth equation and are represented as mean ± S.E.M. (n=3).

### Protein export is unimpaired in PfHsp70x-KO parasites

Using PfHsp70x-KO parasites, we next tested the hypothesis that the chaperone is required for export of virulence-associated proteins. Using immunofluorescence, we examined the export of the same proteins assayed with PfHsp70x-*glmS* parasites: PfFIKK4.2, KAHRP, and MAHRP1 (31–33). Consistent with our observations using PfHsp70x-*glmS*, *pfhsp70x* knockout did not interrupt export of these proteins (**Fig. 5**). These data show that the loss of PfHsp70x does not impede the parasite’s ability to export virulence-associated proteins to the host cell.

**Fig. 5.**
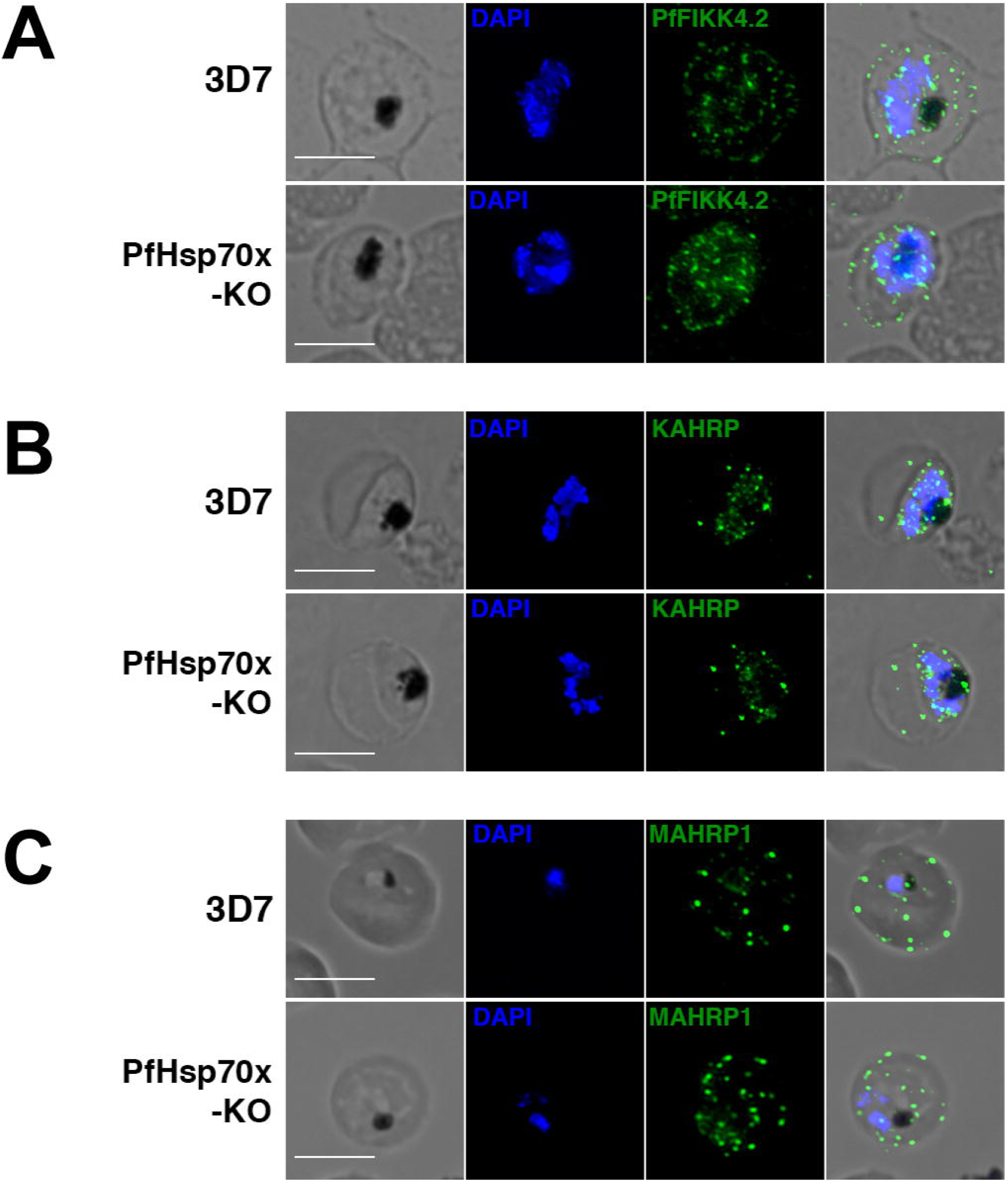
PfHsp70x knockout does not inhibit export of virulence-associated proteins. Asynchronous 3D7 and PfHsp70x-KO parasites were fixed with acetone (PfFIKK4.2 and MAHRP1) or paraformaldehyde (KAHRP) and stained with antibodies against (A) PfFIKK4.2, (B) KAHRP, or (C) MAHRP1. DAPI used to mark parasite cell nucleus. Images from left to right are phase, DAPI (blue), anti-exported protein (green), and fluorescence and phase merge. Scale bar represents 5μm. Representative images shown.

### Export of antigenic proteins to the host RBC is unaffected in PfHsp70x mutants

PfHsp70x was shown to interact with the antigenically variant protein PfEMP1, and recent data that identified proteins that interact with PfEMP1 confirms these results (1). Therefore, we wanted to test the how the export of PfEMP1 is affected in our mutants. Utilizing immunofluorescence microscopy, we determined the localization of PfEMP1 in 3D7 and PfHsp70x-KO parasites (**Fig. 6**). Our data show that knockout of PfHsp70x does not prevent export of PfEMP1 to the host cell (**Fig. 6**). Next, we observed the export of PfEMP1 in our PfHsp70x conditional mutants. Our data show that PfEMP1 is exported equally well in both PfHsp70x-*M9* and PfHsp70x-*glmS* parasites under knockdown conditions (**Fig. 7**). We quantified the amount of PfHsp70x-HA, as well as the amount of exported PfEMP1, in these mutants and found no difference in regards to PfEMP1, despite achieving significant reduction of PfHsp70x in the *glmS* parasite line (**Fig. 7A, B**). Because MAHRP1 has been implicated in the trafficking of PfEMP1, we also quantified the export of MAHRP1 in the PfHsp70x conditional mutants, and we found that knockdown of PfHsp70x does not affect MAHRP1 export (**Fig. 7C**).

**Fig. 6.**
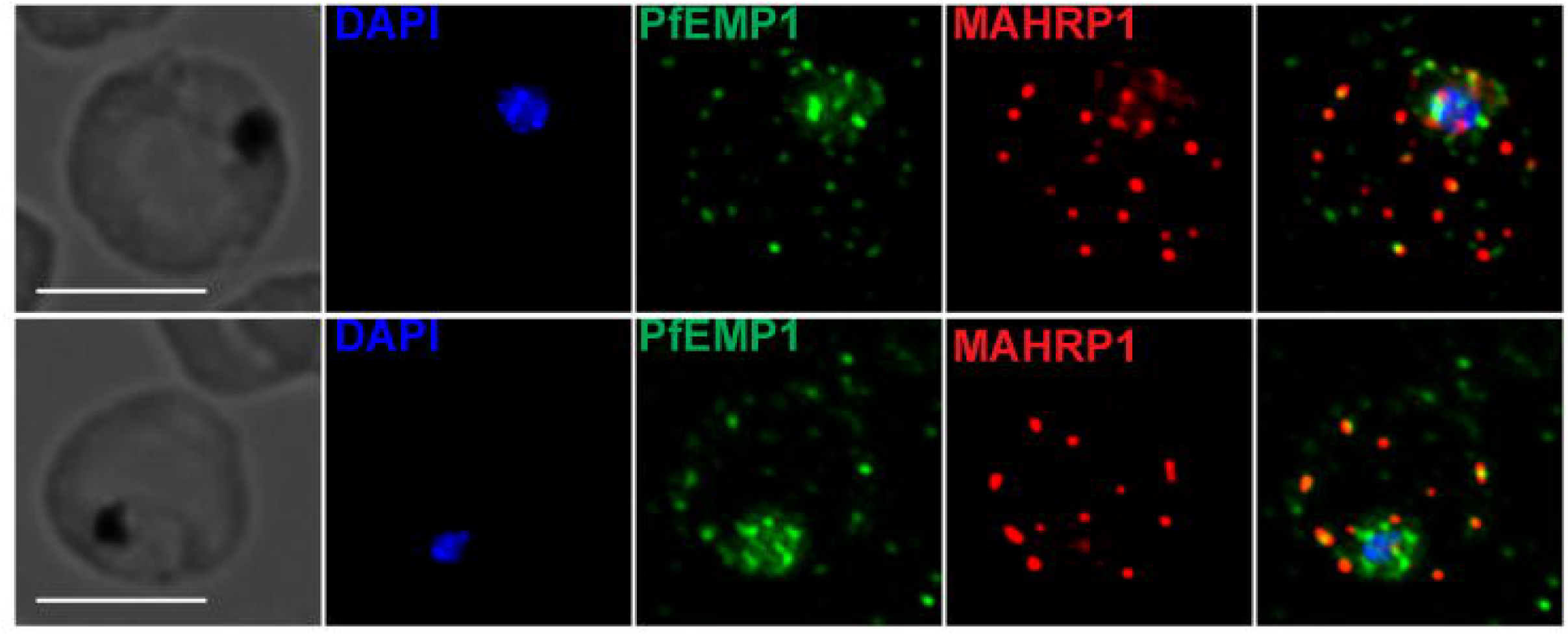
PfHsp70x knockout does not inhibit export of PfEMP1 to the host cell. Asynchronous 3D7 and PfHsp70x-KO parasites were fixed with acetone and stained with antibodies against the ATS domain of PfEMP1 and MAHRP1. DAPI used to stain parasite cell nucleus. Images from left to right are phase, DAPI (Blue), PfEMP1 (green), MAHRP1 (red), and fluorescence merge. Representative images shown.

**Fig. 7.**
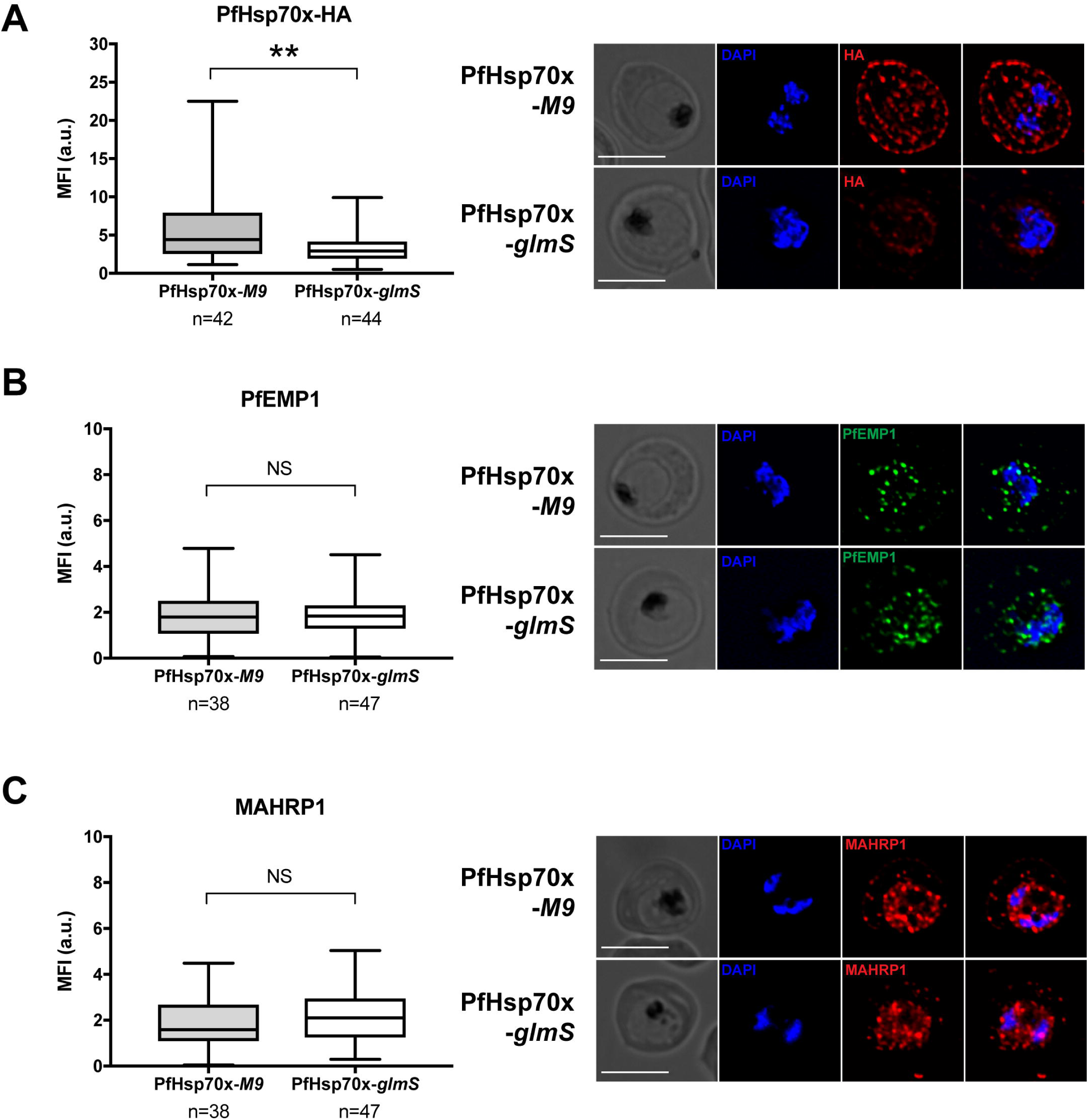
Knockdown of PfHsp70x does not inhibit export of PfEMP1 to the host cell. PfHsp70x-*M9* and PfHsp70x-*glmS* parasites were fixed with acetone and stained with antibodies against (A) HA, (B) PfEMP1, or (C) MAHRP1. DAPI used to mark parasite cell nucleus. Scale bar represents 5μm. Images from left to right are phase, DAPI, anti-HA or –exported protein, and fluorescence merge. Representative images shown. The mean fluorescence intensity (MFI) for each protein was calculated for individual cells and is shown as box-and-whisker plots, with whiskers representing the maximum and minimum MFI. For HA, the MFI was calculated for the entire infected RBC. For PfEMP1 and MAHRP1, MFI was calculated for the exported fraction only. Significance was determined using an unpaired t-test (**, P = 0.01. NS = not significant).

Next, we sought to investigate if there were any differences in the mutants in the export of antigenic parasite proteins that generate an immune response. We obtained pooled human sera collected from a malaria-endemic region (Kenya) as well as a non-endemic region (USA) (34). Uninfected RBCs, 3D7 parasites, and PfHsp70x-KO parasites were labeled with these sera and observed via flow cytometry (**Fig. 8**). 3D7 and PfHsp70x-KO schizonts were synchronized and grown to the schizont stage, and cultures were brought to matching parasitemia prior to labeling with sera. Our data show that both 3D7 and PfHsp70x-KO parasites are labeled equally well by human sera collected from malaria-endemic regions but not by sera obtained from non-endemic regions, suggesting that the export of antigenic parasite proteins to the host RBC is unaffected by the loss of PfHsp70x (**Fig. 8**).

**Fig. 8.**
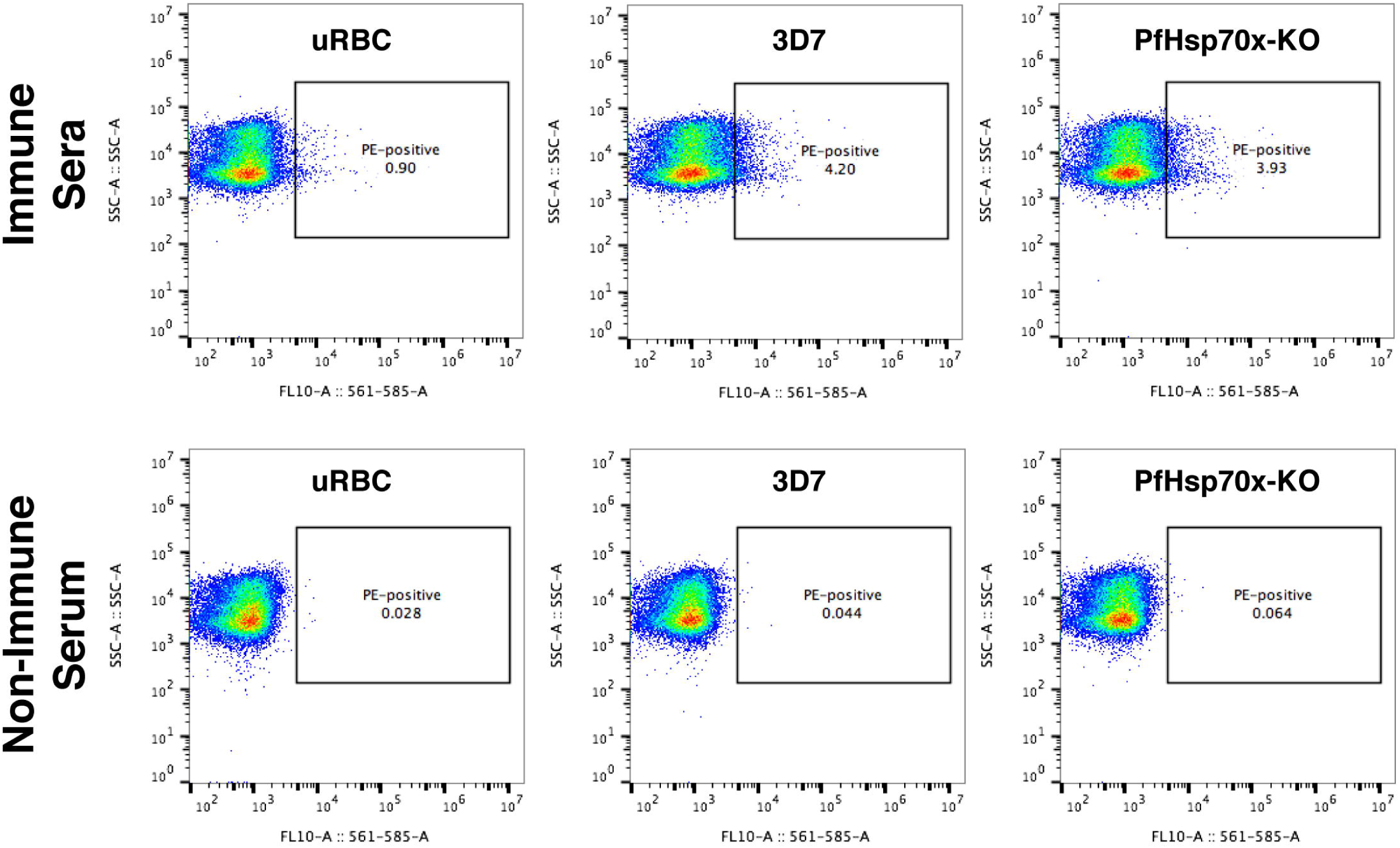
Human immune sera recognizes 3D7 and PfHsp70x-KO parasites. Synchronized 3D7 and PfHsp70x-KO parasites were incubated with either pooled human sera from malaria-endemic Kenya (top panels) or non-immune human serum from the United States. Recognition by the serum was determined using a PE-conjugated anti-human-IgG antibody and flow cytometry. Also assayed were uninfected red blood cells (uRBCs). Side-scatter shown on the Y axis, and PE fluorescence on the X axis.

## Discussion

While this work was under review (and also available on the bioRχiv preprint server), another study was published showing that knockout of PfHsp70x did not affect parasite growth (35). In agreement with these data, our data also demonstrate that PfHsp70x is not required for intraerythrocytic growth, even though PfHsp70x is the only parasite-encoded Hsp70 that is exported to the RBC (**Fig. S2A, Fig. 2A,B,C, Fig.3D and Fig.S3D**). Using two different genetic approaches we demonstrate that the export of several parasite effectors are unaffected by the loss of PfHsp70x (**Fig. 3**, **Fig. 5–8**). In the case of PfEMP1, the newly published work suggests that knockout of PfHsp70x led to delays in its export and minor loss in cytoadherence, suggesting a role for PfHsp70x in parasite virulence (35). In this case, the data show that PfHsp70x knockout parasites over-express some exported proteins (35). This suggests that there may be compensatory mechanisms that are activated when PfHsp70x is knocked out and therefore lead to minor, if any, changes in the export of parasite virulence (35). However, this interpretation is clouded by the lack of a conditional mutant for PfHsp70x, which cannot compensate for the loss of PfHsp70x. The data described in this study show that in both PfHsp70-KO and PfHsp70x-*glmS* mutants, export of parasite virulence factors is not affected (**Fig. 3, Fig. 5–8**). We specifically tested the export of the antigenically variant protein, PfEMP1, which is responsible for cytoadherence, and observed that the export of PfEMP1 was unaffected in either the knockout or the conditional mutants of PfHsp70x (**Fig. 6–8**). Therefore, our data suggest a slightly different, though not mutually exclusive, model than the one proposed in Charnaud *et al*. PfHsp70x is not the only Hsp70 found in infected RBCs. Several human chaperones, including Hsp70, are present in the erythrocyte cytoplasm (36). Thus, the role played by PfHsp70x in the parasite’s biology could be redundant with the human Hsp70 that is already present in the host cell. In fact, infection with *P. falciparum* affects the normal localization of the human Hsp70, as the protein is soluble in non-parasitized RBCs but is found in detergent-resistant fractions following infection (37). Another paper published while this work was under review identified several interacting partners of PfEMP1 using thorough proteomic and genetic data (38). They identified several human chaperones, specifically from the TRiC chaperonin complex, to interact with PfEMP1. Together with our data, this suggests a model wherein PfEMP1 export is aided both by PfHsp70x and by human chaperones present in the host RBCs. This further suggests that loss of either one of them may not be enough to derail the export of parasite virulence proteins to the host RBC. The methods used here to investigate the function of PfHsp70x, knockdown and complete genomic knockout, are more challenging to use for human chaperones such as Hsp70 or the TRiC chaperonin complex. The mature RBC cannot be genetically manipulated, and knockdown of human Hsp70 in hematopoietic stem cells abrogates RBC formation (39). Our data demonstrate that pooled human sera collected from malaria-endemic regions are unable to differentiate between wildtype and PfHsp70x-KO parasites, raising the possibility that PfHsp70x may not be required in human infections (**Fig. 8**). However, further detailed analysis of the *pfhsp70x* locus in strains isolated from the field or testing its role in other stages of the parasite life cycle may be informative about the essentiality of PfHsp70x in human infections. Overall, our data demonstrate that PfHsp70x is not required for export of *P. falciparum* effector proteins to the host, is dispensible for asexual growth within human RBCs, and suggest a model where both human chaperones and parasite chaperones act in a redundant manner to ensure export of parasite virulence factors to the host RBCs.

## Materials and Methods

### Plasmid construction

Genomic DNA was isolated from *P. falciparum* using the QIAamp DNA blood kit (QIAGEN). Constructs utilized in this study were confirmed by sequencing. PCR products were inserted into the respective plasmids using the In-Fusion cloning system (Clonetech) or using the SLIC method. Briefly, insert and cut vector were mixed with a T4 DNA polymerase and incubated for 2.5 minutes at room temperature, followed by 10 minutes incubation on ice and then transformed into bacteria. For generation of plasmid PfHsp70x-HADB, a 1-kb homologous sequence from the 3’-end of the *pfhsp70x* gene (not including the stop codon) was amplified by PCR using primers 5’-CACTATAGAACTCGAGGTGAAAAAGCTAAACGTGTATTATCATCATCCGCACAAGC-3’ and 5’-CGTATGGGTACCTAGGATTTACTTCTTCAACGGTTGGTCCATTATTTTGTGC-3’ and was inserted into pHADB (18) using restriction sites XhoI and AvrII (New England Biolabs).

For the generation of the *glmS* conditional mutants three plasmids were used; 1) pUF1-Cas9 (from J. J. Lopez-Rubio) was used to drive cas9 expression (30). 2) pMK-U6 was used to drive expression of the RNA guide. For this purpose, pL6 plasmid (from J. J. Lopez-Rubio, (30)) was digested with NotI and NcoI (New England Biolabs) and the fragment that contained the U6 RNA expression cassette was blunted and re-ligated to form the pMK-U6 plasmid. The guideRNA, oligos 5’-TAAGTATATAATATTTGCATTATTGTTGTATATTTGTTTTAGAGCTAGAA-3’ and 5’-TTCTAGCTCTAAAACAAATATACAACAATAATGCAAATATTATATACTTA-3’ were annealed and cloned into the RNA module in MK-U6 as previously described (30). Briefly, pMK-U6 was digested with BtgZI (New England Biolabs) and annealed oligos were inserted using In-Fusion HD Cloning Kit (Clontech). 3) pHA-glmS and pHA-M9 were used as donor DNA templates consisting of two homology regions flanking the HA tag and the *glmS* (or the *M9*) sequences. To generate the pHA-glmS and pHA-M9 plasmids, primers 5’-GAGCTCGCTAGCAAGCTTGCCGGCAAGATCATGTGATTTCTCTTTGTTCAAGGAGTC-3’ and 5’-TCCGCGGAGCGCTACTAGTTACCCATACGATGTTCCAGATTACGCTTACCCATACGATGTTCCAGATTACGCTTACCCATACGATGTTCCAGATTACGCTTAAATGTCCAGAC CTGCAGTAATTATCCCGCCCGAACTAAGCGC-3’ were used to amplify the *glmS* and *M9* sequences from pGFP-glmS and pGFP-M9, respectively (from P.Shaw, (26)). PCR constructs were then inserted into a TOPO cloning vector (ThermoFisher). To allow efficient genomic integration of the pHA-glmS and pHA-M9 donor plasmids, 800bp sequences were used for each homology region. The C-terminus of the *pfhsp70x* coding region was PCR amplified form genomic DNA using primers 5’-AATTCGCCCTTCCGCGGGCTGTACAAGCAGCCATCTTATCAGGTGATCAATCATC-3’ and 5’-ATCGTATGGGTAAGCGCTATTTACTTCTTCAACGGTTGGTCCATTATTTTGTGCTTC-3’ and was inserted into pHA-glmS and pHA-M9 using restriction sites SacII and AfeI (New England Biolabs). The 3’UTR of *pfhsp70x* was pcr amplified from genomic DNA using primers 5’-ATGATCTTGCCGGCAAGCTTACGAAAATATACAACAATAATGCATAAAATAATAATA ATT-3’ and 5’-CCTTGAGCTCGCTAGCGCAATATAAATGGATTATTCCTTTTGTATATAATTTAAAATAAG-3’ and was inserted into pHA-glmS and pHA-M9 (already containing the C-terminus homology region) using restriction sites HindIII and NheI (New England Biolabs).

For the generation of *pfhsp70x*-*ko* parasites two plasmids were used; 1) A cas9 expressing plasmid (as described above), and 2) pL7-PfHsp70x plasmid that is derived from the pL6 plasmid (from J. J. Lopez-Rubio, (30)). pL7-PfHsp70x contained the guide RNA and 800bp homology regions flanking a *hdhfr* gene that confers resistance to WR. The N-terminus of the *pfhsp70x* gene was amplified via PCR from genomic DNA using primers 5’-cggggaggactagtATGAAGACAAAAATTTGTAGTTATATTCATTATATTG-3’ and 5’-acaaaatgcttaagGGAAACATCTTTACCTCCATTTTTTTTTTTAAAATCTTGTAC-3’ and was inserted into pL6 using restriction sites AflII and SpeI (New England Biolabs). The C-terminus of the *pfhsp70x* gene was pcr amplified from genomic DNA using primers 5’-taaatctagaattcTGATCAATCATCAGCTGTCAAAGACTTATTATTATTAGATG-3’ and 5’-ttaccgttccatggTTAATTTACTTCTTCAACGGTTGGTCCATTATTTTGTGCTTC-3’ and was inserted into pL6 (already containing the C-terminus homology region) using restriction sites NcoI and EcoRI (New England Biolabs). In order to insert the guide DNA sequence, oligos 5’-TAAGTATATAATATTGTACAAGCAGCCATCTTATCGTTTTAGAGCTAGAA-3’ and 5’-TTCTAGCTCTAAAACGATAAGATGGCTGCTTGTACAATATTATATACTTA-3’ were annealed and cloned into pL6 as previously described (30). Briefly, pL6 was digested with BtgZI (New England Biolabs) and annealed oligos were inserted using In-Fusion HD Cloning Kit (Clontech).

### Cell culture and transfections

Parasites were cultured in RPMI medium supplemented with Albumax I (Gibco) and transfected as described earlier (40, 41). For generation of *PfHsp70x-DDD* parasites, PfHsp70x-HADB was transfected in duplicates into 3D7-derived parental strain PM1KO which contains a *hDHFR* expression cassette conferring resistance to TMP (42). Selection and drug cycling were performed as described (25) in the presence of 10 μM of TMP (Sigma). Integration was detected after three rounds of drug cycling with blasticidin (Sigma).

For generation of PfHsp70x*-glmS* and PfHsp70x*-M9* parasites, a mix of three plasmids (40 μg of each) was transfected in duplicates into 3D7 parasites. The plasmids mix contained pUF1-Cas9 (from J. J. Lopez-Rubio, (30)) which contains the *DHOD* resistance gene, pMK-U6-PfHsp70x, pHA-glmS-PfHsp70x or pHA-M9-PfHsp70x, which are all marker-free. Drug pressure was applied 48 hours post transfection, using 1μM DSM (43), selecting only for Cas9 expression. Drug was removed from the culturing media once parasites became detectable in the culture, usually around 3 weeks post transfection.

For generation of PfHsp70x-KO parasites, a mix of pUF1-Cas9 (from J. J. Lopez-Rubio, (30)) and pL7-PfHsp70x (50 μg of each plasmid) was transfected in duplicates into 3D7 parasites. Drug pressure was applied 48 hours post transfection, using 2.5nM WR99210 (Sigma), selecting for integration of the drug resistance cassette into the *pfhsp70x* gene.

### Growth assays

For asynchronous growth assays of PfHsp70x-DDD lines, parasites were washed twice and incubated without TMP. For asynchronous growth assays of PfHsp70x*-glmS* and PfHsp70x*-M9 parasites*, 5 or 10 mM GlcN (Sigma) were added to the growth media. Asynchronous growth assays of PfHsp70x-KO parasites were performed in media containing WR99210. Parasitemia was monitored every 24 hours via flow cytometry. For flow cytometry, aliquots of parasite cultures (5 μl) were stained with 1.5 mg/ml Acridine Orange (Molecular Probes) in PBS. The fluorescence profiles of infected erythrocytes were measured by flow cytometry on a CyAn ADP (Beckman Coulter) or CytoFLEX (Beckman Coulter) and analyzed by FlowJo software (Treestar, Inc.). Whenever required, parasites were sub-cultured to avoid high parasite density and relative parasitemia at each time point was back-calculated based on actual parasitemia multiplied by the relevant dilution factors. 100% parasitemia was determined as the highest relative parasitemia and was used to normalize parasite growth. Data were fit to exponential growth equations using Prism (GraphPad Software, Inc.)

### Southern blotting

Southern blots were performed with genomic DNA isolated using the Qiagen Blood and Cell Culture kit. 10 μg of DNA was digested overnight with NcoI/XmnI for PfHsp70x-DDD and BamHI/ScaI for PfHsp70x-KO (New England Biolabs). Integrants were screened using biotin-labeled probes against the 3’-end (PfHsp70x-DDD parasites) or 5’-end (PfHsp70x-KO parasites) of the *pfhsp70x* ORF. Southern blot was performed as described earlier (44). The probe was labeled using biotinylated Biotin-16-dUTP (Sigma). The biotinylated probe was detected on blots using IRDye 800CW Streptavidin conjugated dye (LICOR Biosciences) and was imaged, processed and analyzed using the Odyssey infrared imaging system software (LICOR Biosciences).

### Western blotting

Western blots were performed as described previously (27). Briefly, late-stage parasites were isolated on a percoll gradient (Genesee Scientific). For PfHsp70x-DDD parasites, host RBCs were permeabilized selectively by treatment with ice-cold 0.04% saponin in PBS for 10 min. Supernatants were collected for detection of exported parasites proteins and pellets were collected for detection of proteins with the parasite. For PfHsp70x-KO, PfHsp70x*-glmS* and PfHsp70x*-M9* parasites, whole parasite lysates, including the host RBC, were used to detect protein expression and export. The antibodies used in this study were rat anti-HA (3F10, Roche, 1:3000), rabbit anti-PfEF1α (from D. Goldberg, 1: 2000), mouse anti-Plasmepsin V (From D. Goldberg, 1:400), rabbit anti-PfHsp70x (From J. Przyborski, 1:1000). The secondary antibodies that were used are IRDye 680CW goat anti-rabbit IgG and IRDye 800CW goat anti-mouse IgG (LICOR Biosciences, 1:20,000). The western blot images were processed and analyzed using the Odyssey infrared imaging system software (LICOR Biosciences).

### Microscopy and image processing

For detection of HA-tags, PfHsp70x, PfFIKK4.2, and MAHRP1, cells were smeared on a slide and acetone-fixed. For KAHRP detection, cells were fixed with paraformaldehyde and glutaraldehyde. PfHsp70x-HA was detected using rat anti-HA antibody (clone 3F10, Roche, 1:100). MAHRP1 was detected using rabbit anti-MAHRP1 (from Hans-Peter Beck, 1:500). PfFIKK4.2 and KAHRP were detected using mouse anti-PfFIKK4.2 (1:1000) and mouse anti-KAHRP (1:1000 and 1:500, respectively. Both antibodies acquired from David Cavanagh and EMRR). PfEMP1 was detected using mouse-anti-ATS (1B/98-6H1-1, 1:100, Alan Cowman). Secondary antibodies used were anti rat-AlexaFluor 488 or 594, anti rabbit-AlexaFluor 488, and anti mouse-AlexaFluor 488 (Life Technologies, 1:100). Cells were mounted on ProLong Diamond with DAPI (Invitrogen) and were imaged using DeltaVision II microscope system with an Olympus IX-71 inverted microscope using a 100X objective. Image processing, analysis and display were preformed using SoftWorx and Adobe Photoshop. Adjustments to brightness and contrast were made for display purposes. For quantification of PfHsp70x-HA fluorescence, PfEMP1 export, and MAHRP1 export, PfHsp70x-*glmS* and –*M9* parasites were grown in the presence of 7.5 mM GlcN for 72 hours, then fixed and stained with anti-HA, anti-ATS, and anti-MAHRP1 as described above. Cells were imaged as described above. The mean fluorescence intensity (MFI) for each protein was calculated as described (10). Briefly, ImageJ was used to calculate MFI for the whole infected RBC (PfHsp70x) or the infected RBC minus the parasite in order to quantify the exported fraction (PfEMP1and MAHRP1). DIC images were used to exclude the parasite from analysis when calculating the MFI of the PfEMP1 and MAHRP1 exported fraction. Data were plotted using Prism (GraphPad Software, Inc.).

### Human Sera Staining

3D7 and PfHsp70x-KO parasites were synchronized to the ring stage by incubating infected RBCs with 5% D-sorbitol (Amresco, Inc.) for 10 minutes at 37 degrees Celsius. Parasites were washed 3 times with culture medium, then allowed to proceed through the lifecycle to the schizont stage. The cultures were incubated 1:10 with either pooled immune sera from Kenya or non-immune serum from the United States for 30 minutes at 37 degrees Celsius, shaking on an orbital shaker at 880 rpm. The serum was washed from the parasites three times with culture medium, and goat-anti-human IgG Fc conjugated to PE was added to the parasites (1:500, Fisher Scientific, 50-112-8944). The secondary antibody was incubated with the parasites for 30 minutes at 37 degrees Celsius, shaking. Parasites were washed 3 times with culture medium, resuspended in PBS, and fluorescence was measured with a flow cytometer (CytoFLEX, Beckman Coulter) and data analyzed using FlowJo software (Treestar, Inc.). Immune serum samples were collected as described, and all samples have been de-identified (34, 45).

## Acknowledgments

We thank Julie Nelson at the Center for Tropical and Emerging Global Diseases Cytometry Shared Resource Laboratory; Muthugapatti Kandasamy at the University of Georgia Biomedical Microscopy Core; Heather M. Bishop for technical assistance; Jose-Juan Lopez-Rubio for sharing the pUF1-Cas9 and pL6 plasmids; and Dan Goldberg (for anti-Plasmepsin V and anti-EF1α), Hans-Peter Beck (for anti-MAHRP), Jude Przyborski (for anti-PfHsp70x), Alan Cowman (for anti-PfEMP1), and David Cavanagh and EMRR (for anti-FIKK4.2 and anti-KAHRP). This work was supported by grants from the March of Dimes Foundation (Basil O’Connor Starter Scholar Research Award), and the US National Institutes of Health (R00AI099156) to V.M, and (T32 AI060546) to M.A.F.

---

**Fig. S1. Generating PfHsp70x-DDD parasites.** (A) Mechanism of PfHsp70x-DDD conditional inhibition. The *pfhsp70x* locus was modified to contain a triple hemagglutinin (HA) tag and a DHFR-based destabilization domain (DDD). In the presence of trimethoprim (TMP) the DDD is stable and the chaperone is active. Upon TMP removal the chaperone binds the DDD intra-molecularly and cannot interact with client proteins, inhibiting normal activity. (B) Single crossover homologous recombination enables the integration of the plasmid into the 3’ end of the *pfhsp70x* gene (upper panel). Southern blot analysis of genomic DNA (bottom panel) isolated from parasite lines indicated above the lanes. The genomic DNA was digested with AccI. Bands expected from integration of the plasmid into the 3’ end of the *pfhsp70x* gene were observed in two independent transfections. A single band indicative of the parental allele was observed for the parental strain and it was absent in the integrant parasites. (C) PfHsp70x-DDD parasites were incubated without TMP, and Schizont stage parasites were purified on a percoll gradient. Host cell lysates together with exported proteins were isolated using 0.04% cold saponin and were then collected from the supernatant (S). Parasites cells with all non-exported proteins were collected from the pellet (P). Using western blot analysis, the two fractions were analyzed and probed for PfHsp70x expression and export. The membrane was probed with antibodies against HA (top) and Plasmepsin V (loading control, bottom). The protein marker sizes that co-migrated with the probed protein are shown on the left.

**Fig. S2. TMP removal does not affect parasite growth and PfHsp70x localization.** (A) Asynchronous PfHsp70x-DDD parasites were grown with or without 10μM TMP and parasitemia was monitored every 24 hours over 5 days. Data are fit to an exponential growth equation and are represented as mean ± S.E.M. Experiments were done 3 times and biological replicates are shown. (B) Immunofluorescence imaging of acetone fixed PfHsp70x-DDD parasites stained with anti-HA (red) and DAPI (blue). Images from left to right are anti-HA (red), DAPI (blue), fluorescence merge, and phase. Scale bar, 5μm.

**Fig. S3. Heat shock does not inhibit the growth of PfHsp70x-KO parasites**. 3D7 and PfHsp70x-KO clones A7 and B3 were subjected to 40 degree Celsius heat shock for 4 hours, and parasitemia was measured every 24 hours using flow cytometry. Data are fit to an exponential growth equation and are represented as mean ± S.E.M. (n=3).

## References

1. Külzer S, Charnaud S, Dagan T, Riedel J, Mandal P, Pesce ER, Blatch GL, Crabb BS, Gilson PR, Przyborski JM. 2012. Plasmodium falciparum-encoded exported hsp70/hsp40 chaperone/co-chaperone complexes within the host erythrocyte. Cell Microbiol 14:1784–1795.

2. WHO. 2016. World Malaria ReportWorld Health Organization.

3. Cowman AF, Healer J, Marapana D, Marsh K. 2015. Malaria: Biology and Disease. Cell 167:610–624.

4. Nilsson SK, Childs LM, Buckee C, Marti M. 2015. Targeting Human Transmission Biology for Malaria Elimination. PLoS Pathog 11:e1004871.

5. Desai SA. 2014. Why do malaria parasites increase host erythrocyte permeability? Trends Parasitol 30:151–159.

6. Maier AG, Cooke BM, Cowman AF, Tilley L. 2009. Malaria parasite proteins that remodel the host erythrocyte. Nat Rev Microbiol 7:341–354.

7. Marti M, Good RT, Rug M, Knuepfer E, Cowman AF. 2004. Targeting Malaria Virulence and Remodeling Proteins to the Host Erythrocyte. Science 306: 1930–1933.

8. Hiller NL, Bhattacharjee S, Ooij C Van, Liolios K, Harrison T, Lopez-estran C, Haldar K. 2004. A Host-Targeting Signal in Virulence Proteins Reveals a Secretome in Malarial Infection. Science 306:1934–1937.

9. Klemba M, Goldberg DE. 2005. Characterization of plasmepsin V, a membrane-bound aspartic protease homolog in the endoplasmic reticulum of Plasmodium falciparum. Mol Biochem Parasitol 143:183–191.

10. Russo I, Babbitt S, Muralidharan V, Butler T, Oksman A, Goldberg DE. 2010. Plasmepsin V licenses Plasmodium proteins for export into the host erythrocyte. Nature 463:632–6.

11. Boddey J a, Hodder AN, Günther S, Gilson PR, Patsiouras H, Kapp E a, Pearce JA, de Koning-Ward TF, Simpson RJ, Crabb BS, Cowman AF. 2010. An aspartyl protease directs malaria effector proteins to the host cell. Nature 463:627–631.

12. Haase S, Herrmann S, Grüring C, Heiber A, Jansen PW, Langer C, Treeck M, Cabrera A, Bruns C, Struck NS, Kono M, Engelberg K, Ruch U, Stunnenberg HG, Gilberger TW, Spielmann T. 2009. Sequence requirements for the export of the Plasmodium falciparum Maurer’s clefts protein REX2. Mol Microbiol 71:1003–1017.

13. Gruring C, Heiber A, Kruse F, Flemming S, Franci G, Colombo SF, Fasana E, Schoeler H, Borgese N, Stunnenberg HG, Przyborski JM, Gilberger TW, Spielmann T. 2012. Uncovering common principles in protein export of malaria parasites. Cell Host Microbe 12:717–729.

14. de Koning-Ward TF, Gilson PR, Boddey JA, Rug M, Smith BJ, Papenfuss AT, Sanders PR, Lundie RJ, Maier AG, Cowman AF, Crabb BS. 2009. A newly discovered protein export machine in malaria parasites. Nature 459:945–9.

15. Gehde N, Hinrichs C, Montilla I, Charpian S, Lingelbach K, Przyborski JM. 2009. Protein unfolding is an essential requirement for transport across the parasitophorous vacuolar membrane of Plasmodium falciparum. Mol Microbiol 71:613–628.

16. Riglar DT, Rogers KL, Hanssen E, Turnbull L, Bullen HE, Charnaud SC, Przyborski J, Gilson PR, Whitchurch CB, Crabb BS, Baum J, Cowman AF. 2013. Spatial association with PTEX complexes defines regions for effector export into Plasmodium falciparum-infected erythrocytes. Nat Commun 4:1415.

17. Boddey JA. 2014. Inhibition of Plasmepsin V activity demonstrates its essential role in protein export, PfEMP1 display, and survival of malaria parasites. PLoS Biol 12:e1001897.

18. Hodder AN, Sleebs BE, Czabotar PE, Gazdik M, Xu Y, O’Neill MT, Lopaticki S, Nebl T, Triglia T, Smith BJ, Lowes K, Boddey JA, Cowman AF. 2015. Structural basis for plasmepsin V inhibition that blocks export of malaria proteins to human erythrocytes. Nat Struct Mol Biol 22:590–596.

19. Beck JR, Muralidharan V, Oksman A, Goldberg DE. 2014. PTEX component HSP101 mediates export of diverse malaria effectors into host erythrocytes. Nature 511:592–5.

20. Elsworth B, Matthews K, Nie CQ, Kalanon M, Charnaud SC, Sanders PR, Chisholm SA, Counihan NA, Shaw PJ, Pino P, Chan J-A, Azevedo MF, Rogerson SJ, Beeson JG, Crabb BS, Gilson PR, de Koning-Ward TF. 2014. PTEX is an essential nexus for protein export in malaria parasites. Nature 511:587–91.

21. Maier AG, Rug M, O’Neill MT, Brown M, Chakravorty S, Szestak T, Chesson J, Wu Y, Hughes K, Coppel RL, Newbold C, Beeson JG, Craig A, Crabb BS, Cowman AF. 2008. Exported Proteins Required for Virulence and Rigidity of Plasmodium falciparum-Infected Human Erythrocytes. Cell 134:48–61.

22. Rhiel M, Bittl V, Tribensky A, Charnaud SC, Strecker M, Müller S, Lanzer M, Sanchez C, Schaeffer-Reiss C, Westermann B, Crabb BS, Gilson PR, Külzer S, Przyborski JM. 2016. Trafficking of the exported P. falciparum chaperone PfHsp70x. Sci Rep 6:36174.

23. Mesen-Ramirez P, Reinsch F, Blancke Soares A, Bergmann B, Ullrich AK, Tenzer S, Spielmann T. 2016. Stable Translocation Intermediates Jam Global Protein Export in Plasmodium falciparum Parasites and Link the PTEX Component EXP2 with Translocation Activity. PLoS Pathog 12:1–28.

24. Elsworth B, Sanders PR, Nebl T, Batinovic S, Kalanon M, Nie CQ, Charnaud SC, Bullen HE, de Koning Ward TF, Tilley L, Crabb BS, Gilson PR. 2016. Proteomic analysis reveals novel proteins associated with the Plasmodium protein exporter PTEX and a loss of complex stability upon truncation of the core PTEX component, PTEX150. Cell Microbiol.

25. Muralidharan V, Oksman A, Pal P, Lindquist S, Goldberg DE. 2012. Plasmodium falciparum heat shock protein 110 stabilizes the asparagine repeat-rich parasite proteome during malarial fevers. Nat Commun 3:1310.

26. Prommana P, Uthaipibull C, Wongsombat C, Kamchonwongpaisan S, Yuthavong Y, Knuepfer E, Holder AA, Shaw PJ. 2013. Inducible Knockdown of Plasmodium Gene Expression Using the glmS Ribozyme. PLoS One 8:1–10.

27. Muralidharan V, Oksman A, Iwamoto M, Wandless TJ, Goldberg DE. 2011. Asparagine repeat function in a Plasmodium falciparum protein assessed via a regulatable fluorescent affinity tag. Proc Natl Acad Sci U S A 108:4411–6.

28. Pei Y, Miller JL, Lindner SE, Vaughan AM, Torii M, Kappe SHI. 2013. Plasmodium yoelii inhibitor of cysteine proteases is exported to exomembrane structures and interacts with yoelipain-2 during asexual blood-stage development. Cell Microbiol 15:1508–1526.

29. Nacer A, Claes A, Roberts A, Scheidig-Benatar C, Sakamoto H, Ghorbal M, Lopez-Rubio J-J, Mattei D. 2015. Discovery of a novel and conserved Plasmodium falciparum exported protein that is important for adhesion of PfEMP1 at the surface of infected erythrocytes. Cell Microbiol.

30. Ghorbal M, Gorman M, Macpherson CR, Martins RM, Scherf A, Lopez-Rubio J-J. 2014. Genome editing in the human malaria parasite Plasmodium falciparum using the CRISPR-Cas9 system. Nat Biotechnol 32:819–21.

31. Kats LM, Fernandez KM, Glenister FK, Herrmann S, Buckingham DW, Siddiqui G, Sharma L, Bamert R, Lucet I, Guillotte M, Mercereau-Puijalon O, Cooke BM. 2014. An exported kinase (FIKK4.2) that mediates virulence-associated changes in Plasmodium falciparum-infected red blood cells. Int J Parasitol 44:319–328.

32. Watermeyer JM, Hale VL, Hackett F, Clare DK, Cutts EE, Vakonakis I, Fleck RA, Blackman MJ, Saibil HR. 2016. A spiral scaffold underlies cytoadherent knobs in Plasmodium falciparum-infected erythrocytes. Blood 127:343–51.

33. Spycher C, Rug M, Pachlatko E, Hanssen E, Ferguson D, Cowman AF, Tilley L, Beck HP. 2008. The Maurer’s cleft protein MAHRP1 is essential for trafficking of PfEMP1 to the surface of Plasmodium falciparum-infected erythrocytes. Mol Microbiol 68:1300–1314.

34. Perrault S, Hajek J, Zhong K, Owino S, Sichangi M, Smith G, Ping Shi Y, Moore J, Kain K. 2009. Human Immunodeficiency Virus Co-Infection Increased Placental Parasite Density and Transplacental Malaria Transmission in Western Kenya. Am J Trop Med Hyg 80:119–125.

35. Charnaud; SC, Dixon; MWA, Nie; CQ, Chappell; L, Sanders; PR, Nebl; T, Hanssen; E, Berriman; M, Chan; J-A, Blanch; AJ, Beeson; JG, Rayner; JC, Przyborski; JM, Tilley; L, Crabb; BS, Gilson; PR. 2017. The exported chaperone Hsp70-x supports virulence functions for Plasmodium falciparum blood stage parasites. PLoS One IN PRESS.

36. Pasini EME, Kirkegaard M, Mortensen P, Lutz HU, Thomas AW, Mann M. 2006. In-depth analysis of the membrane and cytosolic proteome of red blood cells. Blood 108:791–801.

37. Banumathy G, Singh V, Tatu U. 2002. Host chaperones are recruited in membrane-bound complexes by Plasmodium falciparum. J Biol Chem 277:3902–3912.

38. Batinovic S, McHugh E, Chisholm SA, Matthews K, Liu B, Dumont L, Charnaud SC, Schneider MP, Gilson PR, de Koning-Ward TF, Dixon MWA, Tilley L. 2017. An exported protein-interacting complex involved in the trafficking of virulence determinants in Plasmodium-infected erythrocytes. Nat Commun 8:16044.

39. Egan ES, Jiang RHY, Moechtar MA, Barteneva NS, Weekes MP, Nobre L V, Gygi SP, Paulo JA, Frantzreb C, Tani Y, Takahashi J, Watanabe S, Goldberg J, Paul AS, Brugnara C, Root DE, Wiegand RC, Doench JG, Duraisingh MT. 2015. A forward genetic screen identifies erythrocyte CD55 as essential for Plasmodium falciparum invasion. Science 348:711–4.

40. Drew ME, Banerjee R, Uffman EW, Gilbertson S, Rosenthal PJ, Goldberg DE. 2008. Plasmodium food vacuole plasmepsins are activated by falcipains. J Biol Chem 283:12870–12876.

41. Russo I, Oksman A, Goldberg DE. 2009. Fatty acid acylation regulates trafficking of the unusual Plasmodium falciparum calpain to the nucleolus. Mol Microbiol 72:229–245.

42. Liu J, Gluzman IY, Drew ME, Goldberg DE. 2005. The role of Plasmodium falciparum food vacuole plasmepsins. J Biol Chem 280:1432–1437.

43. Ganesan SM, Morrisey JM, Ke H, Painter HJ, Laroiya K, Phillips MA, Rathod PK, Mather MW, Vaidya AB. 2011. Yeast dihydroorotate dehydrogenase as a new selectable marker for Plasmodium falciparum transfection. Mol Biochem Parasitol 177:29–34.

44. Klemba M, Gluzman I, Goldberg DE. 2004. A Plasmodium falciparum dipeptidyl aminopeptidase I participates in vacuolar hemoglobin degradation. J Biol Chem 279:43000–43007.

45. Avery JW, Smith GM, Owino SO, Sarr D, Nagy T, Mwalimu S, Matthias J, Kelly LF, Poovassery JS, Middii JD, Abramowsky C, Moore JM. 2012. Maternal malaria induces a procoagulant and antifibrinolytic state that is embryotoxic but responsive to anticoagulant therapy. PLoS One 7:1–15.

